# Through hawks’ eyes: reconstructing a bird’s visual field in flight to study gaze strategy and attention during perching and obstacle avoidance

**DOI:** 10.1101/2021.06.16.446415

**Authors:** Sofía Miñano, Graham K. Taylor

## Abstract

We present a method to analyse visual attention of a bird in flight, that combines motion capture data with renderings from virtual cameras. We applied it to a small subset of a larger dataset of perching and obstacle avoidance manoeuvres, and studied visual field stabilisation and gaze shifts. Our approach allows us to synthesise visual cues available to the bird during flight, such as depth information and optic flow, which can lead to novel insights into the bird’s gaze strategy in flight. This preliminary work demonstrates the method and suggests several new hypotheses to investigate with the full dataset.

## 1. Introduction

Birds of prey rely strongly on vision for executing their flight manoeuvres, and as a result have developed specialised traits in their visual systems [16, 20]. For example, Harris’ hawks (*Parabuteo unicinctus*) have two high-acuity regions in each retina, called foveae, looking frontally and laterally in the visual field [16, 19, 24]. Studies investigating the role of vision in the fast flight manoeuvres of birds typically involve simplified visual environments, such as corridors with striped patterns on the walls or floor [6, 2]. Here, we present a method that allows us to study in detail how birds control their gaze in flight, in a more realistic visual scenario. Our approach combines motion capture experiments on hawks with tools from computer vision, allowing us to synthesise different vision-based inputs that could potentially be available to the bird during flight (Figure 1). Examples include depth maps (distance measurements per pixel), optic flow data (motion patterns between consecutive frames, in pixel space) or semantic maps (which describe for each pixel the object it belongs to). Inputs such as these are often used in bio-inspired computer vision applications for navigation [18, 14, 22, 1], and can provide novel insight into bird behaviour. We demonstrate the method by applying it to a small sample of flights of a Harris’ hawk executing perching and obstacle avoidance manoeuvres, which is part of a larger dataset that will be analysed fully elsewhere.

**Figure 1:**
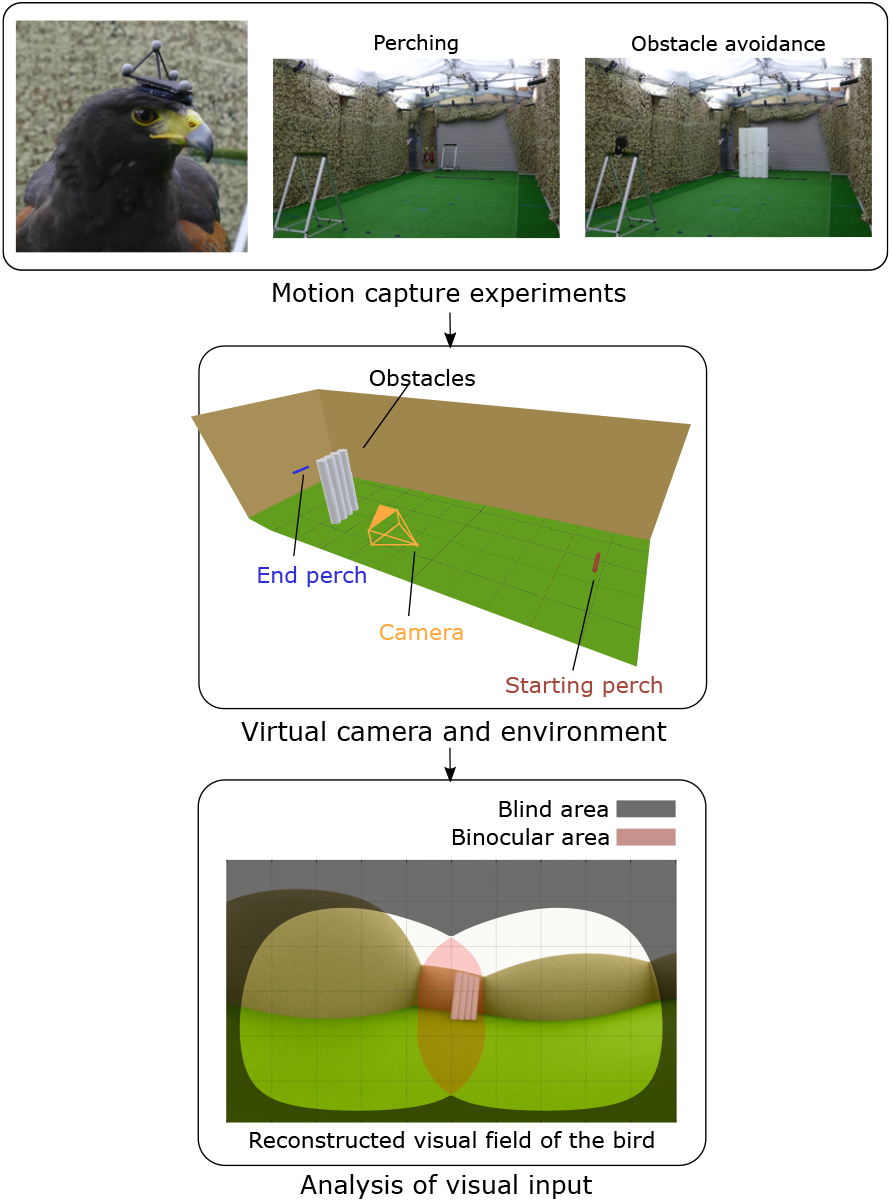
A computer vision method to investigate visual attention of birds in flight. We imported motion capture data from a flying bird into a computer graphics package, to synthetically reconstruct its visual field in flight. With the rendered data we can inspect in detail how the bird stabilises and directs its gaze.

## 2. Methods

We tracked the head movements of a Harris’ hawk executing perching and obstacle avoidance manoeuvres in a motion capture lab. From the data collected across 15 trials, we estimated a coordinate system aligned to the bird’s visual field. We then used a 3D computer graphics package to define a virtual camera that followed the bird’s motions through a simplified geometry of the lab. For two selected trials (one for each type of manoeuvre), we produced RGB renderings, depth maps and semantic maps. We combined these with a map of the bird’s visual field to analyse its gaze stabilisation and gaze shifts in flight. A brief summary of the method is provided here, but for full details, please see the Supplementary Material [15].

### 2.1. Motion capture experiments

To track the bird’s head, we designed a 3D-printed “headpack” with four retroreflective markers (ø6.4 mm), which we fixed to the bird with a Velcro patch glued to the bird’s head (see supplied Supplementary Material [15]). The bird flew in a volume of size 12.0 × 5.3 × 3.3 m with 22 motion capture cameras (Vicon Vantage V16; 200 Hz sampling rate). In the perching trials, the bird flew from a starting perch to an end perch (first leg of the trial), and back (second leg). We randomised the perches’ lateral position (see [15]). In the obstacle avoidance trials, we added four cylindrical styrofoam pillars of 0.3 m diameter and 2 m height, 1 m ahead of the end perch. We fixed large markers (ø14 mm) to the edges of the perches and the tops of the obstacles. We present motion capture data for one bird on one day of experiments (7 perching trials and 8 obstacle avoidance trials). Data collected for three other birds on the same day are included in [15].

We used the software Vicon Nexus 2.8.0 to extract the unlabelled 3D coordinates of the retroreflective markers, and custom MATLAB [13] scripts to label them, handle missing data, and extract the headpack pose (see [15]). To estimate the walls’ location, we used the position and orientation of the motion capture cameras themselves, which were mounted on scaffolding around the walls (see [15]).

### 2.2. Coordinate systems

From the labelled headpack markers, we derived the rotation and translation of a headpack coordinate system relative to the world reference frame. However, this coordinate system is not necessarily relevant for analysing the bird’s visual field, since the headpack is arbitrarily placed on the head. For this purpose we define the *visual coordinate system*: a biologically meaningful reference frame, that allows us to compare results across birds and days, and to previous literature. We additionally define the *trajectory coordinate system*, as one that looks in the forward direction of the head trajectory. We use it for comparison when analysing visual field stabilisation.

To define the *visual coordinate system*, we make use of three assumptions: (1) we consider the bird’s head orientation as a proxy for its eye orientation [11, 5, 23, 12, 10, 7] (a range of eye motion ≤ ±3° is estimated for goshawks, Accipiter gentilis, in [9]); (2) we assume that the bird fixates its gaze on the perch’s centre upon landing [19, 12]; and (3), we assume that the bird keeps its eyes level in flight [4, 23, 25]. We use these assumptions to estimate the bird’s gaze direction 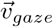 and the normal 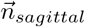 to the head’s symmetry plane (the sagittal plane), which define the axes of the visual coordinate system. We define its x-axis parallel to 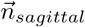 (pointing to the left side of the head) and its y-axis parallel to 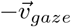.

To estimate the gaze direction unit vector 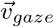, we fitted a line to the trajectories of the landing perch midpoint in the headpack coordinate system (Figure 2). We used data from all 15 trials during the final approach phase, which we defined based on the distance to the landing perch (0.5 to 2.0 m for perching trials, 0.5 to 1.0 m for obstacle trials). Results were similar to those obtained by fitting each trial individually (RMSE = 65.6 mm, 1274 samples, see [15]). To estimate the normal to the sagittal plane 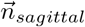, we com-pute the unit vector perpendicular to 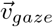 that best approximates the local horizon in the headpack coordinate system during the mid-flight phase. We defined this phase to be when the bird was > 2 m away from either perch and flying at a speed > 2.5 m/s. We identify 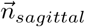 as the solution to the least-squares problem:

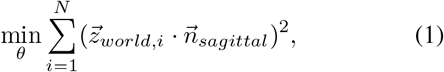

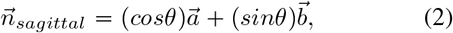

where 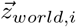 is the world z-axis in the headpack coordinate system (positive opposite to gravity), 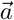 and 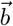 are an arbitrary orthonormal basis of the plane perpendicular to 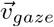, and *θ* is the angle between 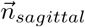 and 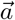. The fit over the 15 trials yields a root-mean square residual of 0.0792 (with *N* = 5056 samples).

**Figure 2:**
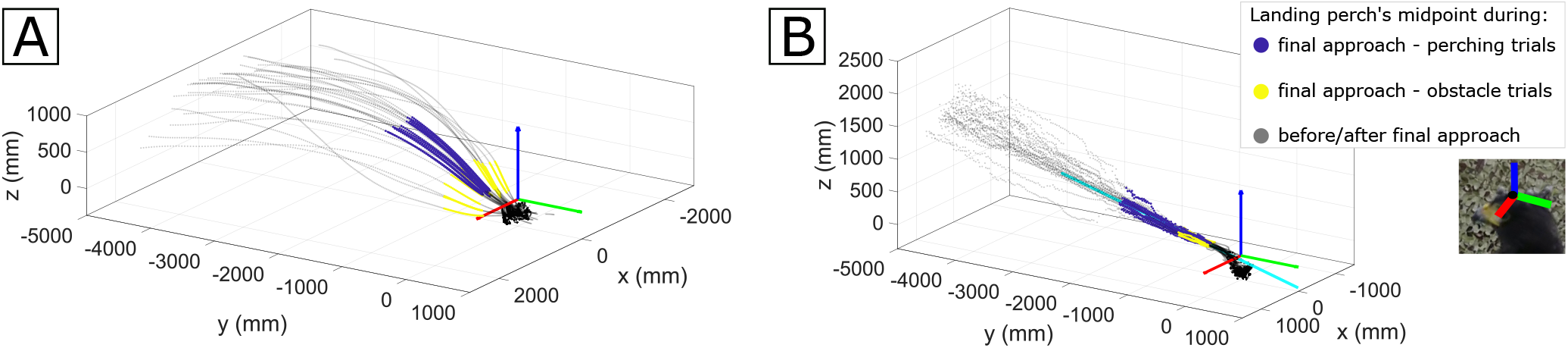
Estimation of gaze direction. The gaze direction 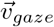 is estimated over 7 perching and 8 obstacle trials (2 legs each). The landing perch’s midpoint (grey and coloured dots) is represented from 5 m away until landing, in two coordinate systems: one that translates with the headpack and is parallel to the world coordinate system (A), and one fixed to the headpack (B). The estimated gaze direction 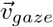 is the orthogonal regression line (cyan) to the samples in the final approach period (plotted blue for perching trials, and yellow for obstacle trials) in the headpack coordinate system (B). Note the straightness of the perch’s midpoint trajectories in the headpack coordinate system in (B), and compare this to the curvature of the lines in (A), which confirms unequivocally that the head pose was being stabilised in relation to the perch. Inset in (B) shows the approximate orientation of the headpack coordinate system relative to the bird’s head.

We define the *trajectory coordinate system* as the axis system with its y-axis parallel to the head’s velocity, and its x-axis parallel to the horizon (in line with the eyes-level assumption, see the Supplementary Material [15]).

The origin of both coordinate systems is defined as the centroid of the headpack markers projected onto the plane defined by the headpack baseplate. We estimate that this origin is within 12 mm of the midpoint between the bird’s eyes in the transverse direction, and < 20 mm in the dorsoventral direction (see the Supplementary Material [15]).

### 2.3. Visual field map

We determined the retinal margins of the bird’s eyes using data from Potier *et al*. [19] (see the Supplementary Material [15]). We identified the direction of maximum binocular overlap with the gaze direction 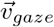 (Fig. S1 in [19]).

### 2.4. Rendering of visual input

We used the Python API in Blender 2.79a [3] to model the simplified geometry of the lab and to define the animation keyframes of a 360° virtual camera. Given that any vergence movements of the eyes were unknown, we modelled the binocular visual field using a monocular camera with equirectangular projection. For the purposes of this paper, we considered a simplified geometry of the environment, given that the aim is to determine gaze strategy in relation to the main elements in the scene. The method allows for the creation of a more detailed model of the scene as required. For the two selected trials, we produced RGB, depth and semantic maps, with a camera following the visual and the trajectory coordinate systems.

## 3. Results

We used the rendered data to investigate the bird’s visual field stabilisation and gaze-shifting behaviour. We analysed the first and second legs of each trial separately (those not in the main text are included in [15]).

To analyse stabilisation, we inspected where in the visual field the landing perch and the obstacles were most frequently seen throughout a flight. Figure 3 shows the results for the first legs of each trial, for a virtual camera moving with the visual coordinate system (top row) and the trajectory coordinate system (bottom row).

**Figure 3:**
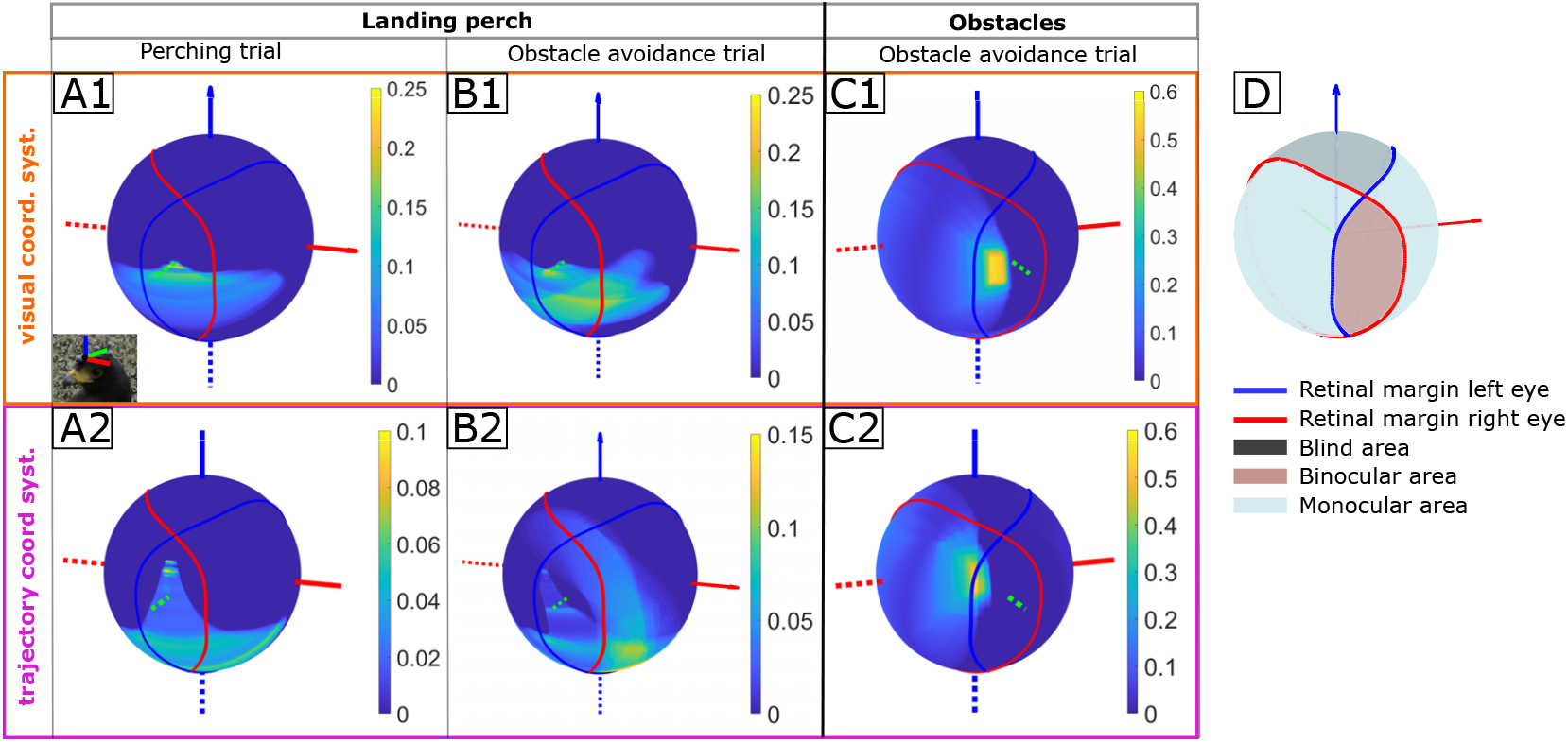
Visual field stabilisation. Heatmaps show the frequency with which an object appears at each point in the visual field, throughout a flight. We computed these heatmaps for the first leg of the selected perching and obstacle avoidance trials. Results are shown for a virtual camera following the visual coordinate system (**A1 to C1**), and the trajectory coordinate system (**A2 to C2**). In the visual coordinate system, objects show considerable stabilisation around the x-axis, and particularly the obstacles’ leftmost edge is strongly aligned with the bird’s sagittal plane (C1). Inset in A1 shows the approximate orientation of the visual coordinate system relative to the bird’s head. Colorbars are scaled based on the rounded-up maximum value observed per heatmap. (**D**) Map of the bird’s visual field, derived from Potier *et al*. [19].

In the visual coordinate system, the perch is stabilised within the binocular area throughout the flight, for both trials (Figure 3, A1, A2, B1 and B2). More evident however, is the stabilisation of the obstacles (Figure 3, C1 and C2): in the visual coordinate system, their leftmost edge is strongly aligned with the bird’s sagittal plane for over 50% of the frames. This is not the case in the trajectory coordinate system, where the obstacles appear most frequently at the boundary of the binocular area.

To investigate where the bird is fixating its gaze and what may motivate a gaze shift, we focused on the second leg of the obstacle avoidance trial. We plotted gaze rays along the trajectory (the gaze vector scaled with the measured depth in that direction), and selected two gaze shifts after the bird had turned around the obstacles, from visual inspection of Figure 4A.

**Figure 4:**
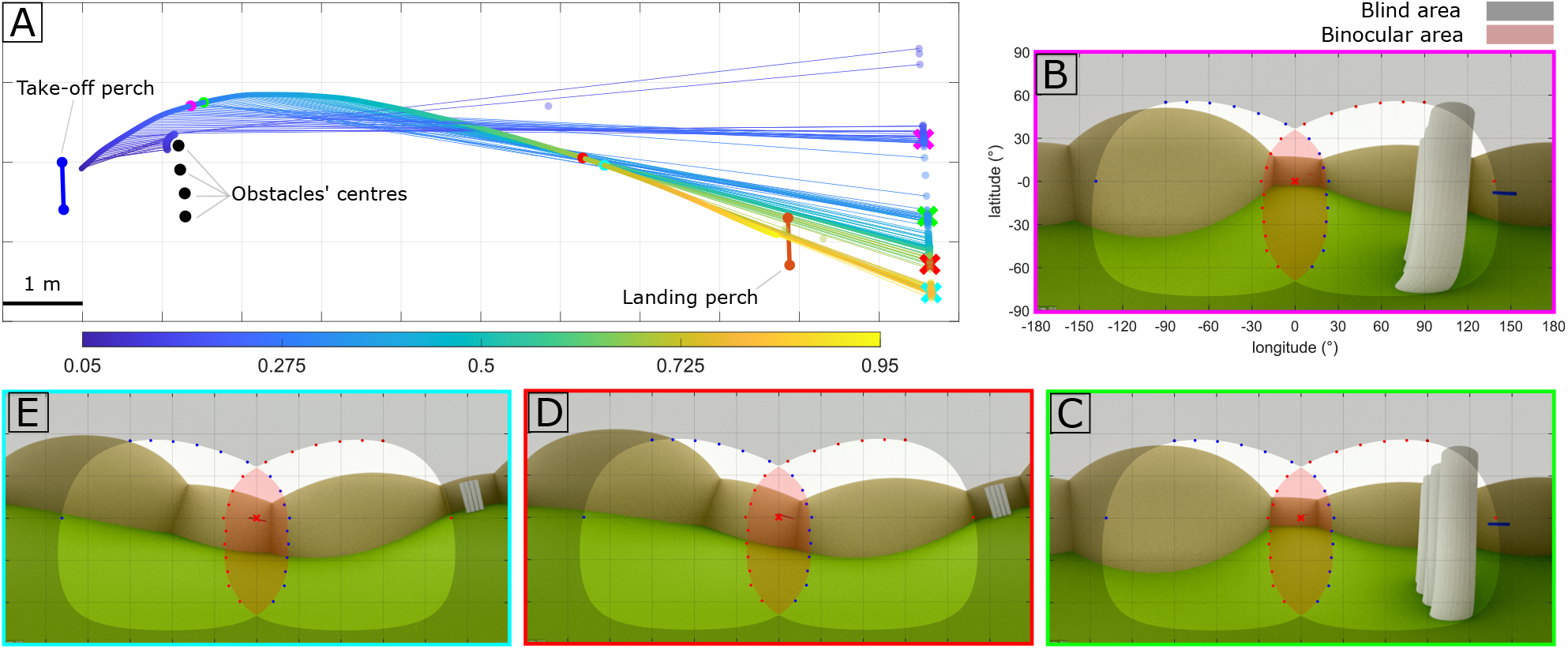
Gaze shifts in an obstacle avoidance flight. **(A)** Top view of the bird’s trajectory on the second leg of the obstacle avoidance trial, with gaze rays shown every other frame. The colormap indicates time through the trajectory (20 first and last frames excluded). The gaze rays’ tips (transparent markers) are plotted for each frame, and marked with crosses at the selected gaze-shifting frames (magenta, green, red, cyan). The selected frames are also marked on the head trajectory as dots with of same colours. **(B-E)** Rendered views at selected frames, with retinal margin data from Potier et al. [19] for the left (blue) and right (red) eyes. The gaze direction is marked with a red cross. The gaze shift between (B) and (C) seems to align the gaze direction with the perch’s left edge. The gaze shift from (D) to (E) seems to align with the perch’s centre. None of the frames is within the final approach phase used to estimate 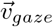. Axes’ labels in C to E omitted for clarity.

The first gaze shift takes 35 ms (magenta to green marker in Figure 4A) and seems to be a reorientation soon after the landing perch is visible, to align with its left edge (Figure 4B and C). This edge seems to be tracked smoothly until approximately the second gaze shift (40 ms, red to cyan marker), when the bird seems to align with the centre of the landing perch (Figure 4D and E). The last frame (cyan) is 50 frames ahead of the final approach phase used to estimate 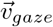 in Section 2.2.

## 4. Conclusions

Our visual field heatmaps agree with previous literature [23], in showing that the bird compensated the wingbeat induced oscillations that would otherwise affect the head pose, to stabilise the perch and obstacles in the visual field. The stabilisation of the obstacles’ edge in the sagittal plane is also in line with similar observations in lovebirds [12] and in humans [21]. Both have been reported to fixate on the edge of objects to avoid (*i.e*., obstacles), and on the centre of objects to aim to (*i.e*., goals). We consider this is a good confirmation that the visual coordinate system estimation is reasonable. This goal-aiming fixation strategy could also explain the gaze shifts identified in Figure 4, where the bird seemed to align first with the perch edge, and then with the perch centre. However, this needs to be confirmed with data from more flights.

In this analysis we have implicitly identified the bird’s gaze direction with that of the forward-facing fovea. However, Harris’ hawks have a lateral-facing fovea as well. With more data, it would be interesting to determine whether both foveae are directed at similar semantic features of the environment in these flights, and whether distance is the main factor determining the use of one or the other, as has been suggested elsewhere [19]. This preliminary study has enabled us to explore the capabilities of the method, and define some specific hypotheses to investigate with the full dataset. A more detailed model of the environment could also benefit the analysis of the bird’s behaviour. We plan to incorporate this next, by making use of detailed 3D maps of the environment, which we have already collected. These maps were generated with the SemanticPaint system described in [8], using an AR smartphone with a depth camera (ASUS ZenFone).

Our method could also inform analyses based on videos from head-mounted cameras. This is a frequent approach in vision research with free-flying birds ([17, 9, 5]). Our method is applicable to a calibrated motion capture environment, but we expect it to be more precise and less impactful on the bird’s behaviour than the use of a head-mounted camera (the headpacks weigh less than 4 g so their attachment is in principle simpler than that of a camera). It could be interesting to compare both approaches, and possibly derive recommendations for head-mounted camera studies.

## 5. Ethics statement

This work received approval from the Animal Welfare and Ethical Review Board of the Department of Zoology, University of Oxford, in accordance with University policy on the use of protected animals for scientific research, permit no. APA/1/5/ZOO/NASPA, and is considered not to pose any significant risk of causing pain, suffering, damage or lasting harm to the animals. No adverse effects were noted during the trials.

## 6. Acknowledgements

We thank our falconers Helen Sanders and Lucy Larkman for animal training and husbandry. We thank Marco Klein Heerenbrink for his assistance with the bird experiments and for helpful suggestions, and Juan Miñano for many useful discussions. We are grateful to Stuart Golodetz for valuable comments on the manuscript, and thank the two anonymous reviewers from the CVPR 2021 workshop CV4Animals for helpful suggestions. This project has received funding from the European Research Council (ERC) under the European Union’s Horizon 2020 research and innovation programme (Grant Agreement No. 682501). SM’s work was supported by funding from the Biotechnology and Biological Sciences Research Council (BBSRC) [grant number BB/M011224/1], via the Intersdisciplinary Bioscience Doctoral Training Partnership.

