## Supplementary Material for "Through hawks’ eyes: reconstructing a bird’s visual field in flight to study gaze strategy and attention during perching and obstacle avoidance"

### Contents

1. Motion capture experiments **1**
2. Postprocessing of motion capture data **2**
3. Visual coordinate system **3**
4. Trajectory coordinate system **4**
5. Distance from the origin of the visual coordinate system to the midpoint between the eyes **5**
6. Visual field map **6**
7. Model of the environment **6**
8. Rendering of visual input **6**
9. Visual field stabilisation **7**
10. Gaze-shifting behaviour **8**
11. Supplementary videos **8**

#### 1. Motion capture experiments

**Motion capture system.** Experiments were conducted in a motion capture lab equipped with 22 infrared motion capture cameras (Vantage V16, Vicon Motion Systems Ltd, Oxford, UK; sampling rate 200 Hz) and four video cameras (Vue, Vicon Motion Systems Ltd, Oxford, UK; sampling rate 100 Hz, only used for reference), covering a volume of size 12.0 × 5.3 × 3.3 m. To track the birds' head movements, we specially designed 3D-printed rigid supports of retroreflective markers, for the birds to wear as a 'headpack'. We used markers of 6.4 mm diameter and attached the headpack to the birds' heads using a Velcro strip glued to their heads (see Figure 1). The birds wore different headpack designs because we used the collected data to validate a custom headpack tracking algorithm, which will be fully described elsewhere.

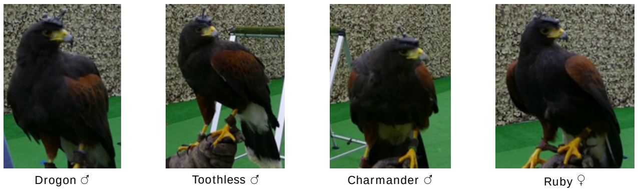


Figure 1: **Birds and headpacks.** We collected motion capture data from four Harris's hawks (*Parabuteo unicinctus*) wearing four headpack designs. The main text focuses on the dataset collected with Drogon.

**Birds.** We worked with four captive-bred adult Harris' hawks, comprising three males (Drogon, Toothless, Charmander) and one female (Ruby), trained to fly between perches. We recorded around 16 flights per bird per day, for a total of two weeks (including the training period for the birds) in November 2020. The data presented in this paper are from flights recorded on one day of experiments, after six days of training. The main text focuses on the trials of one bird (Drogon).

**Experimental setup.** For each recorded trial, the bird would fly from the starting perch to the end perch, and back. Two falconers stood at either end of the room, to handle the bird and provide the food reward. The perches were placed 9 m apart along the longitudinal axis of the room (Figure 2). Their lateral position was randomised for every trial between three stations each. These were centred on the longitudinal axis of the room and distributed with 1 m spacing. For the obstacle avoidance trials, we placed four styrofoam pillars of 2 m height and 0.3 m diameter, 1 m ahead of the end perch (Figure 2). Each obstacle was made up of two white expanded polystyrene cylinders stacked on top of each other and bound together with white duct tape. The four obstacles were pushed together so that there were minimal gaps between them. In the obstacle avoidance trials, we also randomised the side of the end perch from which the falconer would call the bird. To reconstruct the geometry of the environment, we placed larger retroreflective markers (14 mm diameter) on the perches' edges and on the obstacles' tops.

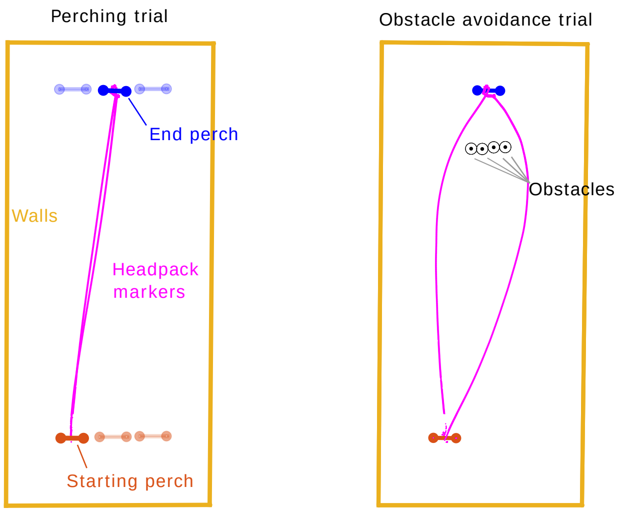


Figure 2: **Schematic of experimental setup (top view).** In the perching trials (left) the bird flew from the starting perch (red) to the end perch (blue) and back. The perches were 9 m apart. Their lateral position was randomised across trials between three stations each, distributed with 1 m spacing (transparent). In the obstacle avoidance trial, we placed a set of four obstacles 1 m ahead of the end perch (black cylinders, centres highlighted with black markers). The reconstructed headpack markers are shown in magenta. The trajectories correspond to the rendered perching trial (left) and obstacle avoidance trial (right). The computed walls position is shown (ochre).

**Experimental procedure.** We calibrated or recalibrated the motion capture system before each trial. The headpack was attached to the head of the bird by the falconer immediately before its set of trials, and removed at the end. For the first half of the trials, we recorded perching manoeuvres. For the second half, we recorded obstacle avoidance manoeuvres. To ensure that the headpack had not moved during the flights, we recorded videos of the birds on the falconer's fist, wearing the headpacks, before their complete set of trials. For all days except two, we also recorded these videos after the bird's complete set of trials. We recorded videos during the trials as well, for reference.

**Dataset.** The motion capture data presented in this paper are from flights recorded on one day of experiments, after six days of training. We reviewed the trials from the selected day and discarded those with invalid take-offs or landings. We considered a take-off or a landing invalid if the bird did not touch the perch with both legs. In the valid trials (those with valid take-offs and landings), we manually determined take-off and landing frames by inspecting the motion capture data and the reference videos. We identified the take-off and landing frames as the point at which the head was lowest when the bird was about to leave the perch or had just landed on it. The number of valid trials per bird for the selected day of experiments is shown in Table 1. The main text focuses on the data collected with Drogon.

Table 1: **Valid trials per bird and manoeuvre type.** A trial is considered valid if both legs have valid take-offs and landings. The main text focuses on the dataset from Drogon.

| Bird | number of perching trials | number of obstacle avoidance trials |
|---|---|---|
| Drogon (♂) | 7 | 8 |
| Toothless (♂) | 4 | 6 |
| Charmander (♂) | 7 | 5 |
| Ruby (♀) | 6 | 8 |

#### 2. Postprocessing of motion capture data

We used the commercial software from the motion capture system (Vicon Nexus 2.8.0) to extract the unlabelled 3D coordinates of the retroreflective markers in time. We wrote custom MATLAB [3] scripts to separate headpack and object markers, label the headpack markers, extract the pose of the headpack in time, and compute the simplified geometry of the environment.

**Headpack transform.** For the few frames where there were not enough headpack markers reconstructed, or where these could not be reliably labelled, we interpolated the headpack's translation using a weighted cubic spline that takes into account the number of markers detected. For the headpack's rotation, we used the SLERP algorithm for quaternions [6], which assumes constant angular velocity. As a verification of the quality of the labelling, we computed the labelled markers in the headpack coordinate system, for the frames in which the headpack transform was defined (i.e., not interpolated). For all the trials across all birds on the selected day, the labelled markers deviated less than 5 mm from their corresponding centroid in the headpack coordinate system.

**Walls.** The motion capture cameras were mounted on a fixed scaffold from which camouflage netting was hung. To minimise sources of noise, we decided not to place additional markers on the netting. Instead, we estimated its location using the position and orientation of the motion capture cameras themselves. These were computed during the calibration, and were the same for all trials per bird. We used the cameras' positions and orientations to estimate points on the scaffolding, taking into account the dimensions of the cameras and their mounts. We then fitted the estimated scaffolding points to a line for each wall section. This provided an estimate of the scaffolding rungs holding the netting and the cameras. We defined each wall as the plane perpendicular to the floor that contained the corresponding scaffolding rung, from which the camouflage netting was hung.

Figure 3 shows the estimated walls and the motion cameras for the Drogon trials. The mean angle at the walls' corners was 90° (with standard deviation $\sigma = 0.5$). We set the ceiling at the mean height of the estimated scaffolding points (3.25 m).

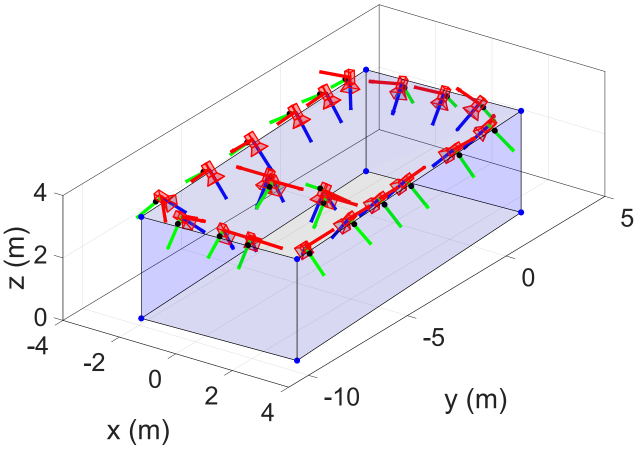


Figure 3: **Estimated location of the walls.** We estimated the location of the walls from the motion capture cameras' positions and orientations. Cameras are shown in red. Each camera's coordinate system is represented with red, green and blue axes (for x, y, z axes respectively). Using estimates of the dimensions of the cameras and their mounts, we estimated points in the scaffolding (black markers). We then used the scaffolding points to the define the walls' planes (blue transparent planes). The corners of the volume are highlighted with blue markers.

#### 3. Visual coordinate system

**Assumptions for estimating the visual coordinate system.** To estimate the visual coordinate system, we assume that the bird fixates its gaze on the perch midpoint on final approach and that it keeps its eyes level in flight (see Section 2.2 in the main text). These assumptions are based on reports on previous literature [1, 5, 4, 2, 8] and on observations from our experiments (see Figure 4).

**Gaze direction estimate.** If the bird is fixating its gaze on the landing perch midpoint in the final approach to the perch, this point's trajectory would be close to a straight line in the headpack coordinate system. This line would

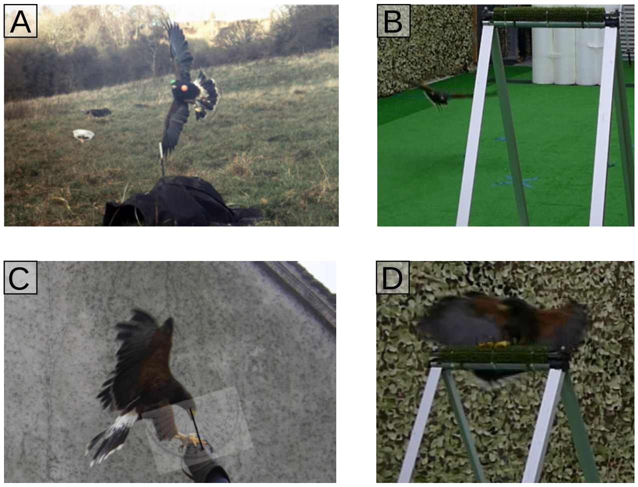


Figure 4: **Perch fixation and eyes-level assumption.** Examples of a bird keeping its eyes level in flight, from literature [1] **(A)** and from our experiments **(B)**. Examples of a bird fixating on the perch upon landing, from literature [4] **(C)** and from our experiments **(D)**.

then be a reasonable estimate of the gaze direction, $\vec{v}_{gaze}$. We define the final approach phase based on the distance between the bird and the perch, 0.5 to 2.0 m for perching trials and 0.5 to 1.0 m for obstacle trials. We used different thresholds depending on the type of trial because we found gaze shifts up to approximately 1 m ahead of the perch in the obstacle avoidance trials (see Figure 5). Note that the obstacles are 1 m ahead of the end perch.

We computed the orthogonal regression line to the trajectories of the perch midpoint in the headpack coordinate system, considering all trials of the same bird and day (i.e., all trials in which the bird had the same headpack placement). We followed the singular-value decomposition approach described in [7]:

$$USV^T = svd(X - \overline{X}), \quad (1)$$

$$\vec{v}_{gaze} = U(:, 1), \quad (2)$$

$$A_{gaze} = \overline{X} \quad (3)$$

where $X$ represents an array of size $3 \times N$ (with $N$ the number of sample points) and $\overline{X}$ is the mean of the sample points (an array of size $3 \times 1$). The singular-value decomposition is represented by the operator $svd()$. The matrices $U$, $S$, $V$ correspond to the matrix containing the left singular vectors in columns, the diagonal matrix containing the singular values, and the matrix containing the right singular vectors in columns, respectively. The estimated gaze direction unit vector $\vec{v}_{gaze}$ is computed as the first column of $U$. $A_{gaze}$ is the centroid of the sample points, and a point in the fitted line. The results of the fits across all trials per bird are summarised in Table 2.

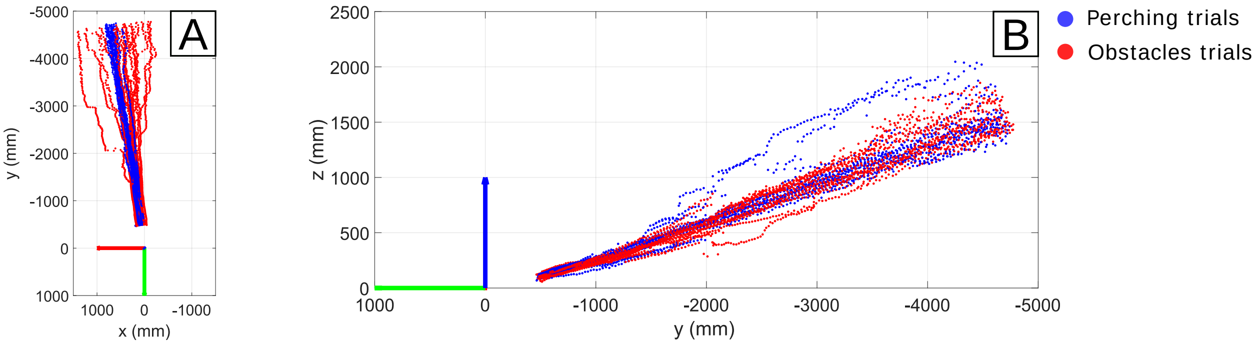


Figure 5: **Trajectories of the landing perch midpoint in the headpack coordinate system.** Trajectories shown for when the bird is 5 m away from the perch until it is 0.5 m from the perch, for all trials of one bird (Drogon). **(A)** Projection onto the *xy* plane. Note the head shifts (in the shape of steps) in the obstacle trials (red). **(B)** Projection onto the *yz* plane.

The root mean square error (RMSE) in Table 2 is computed with the error being the distance between the samples and the fitted line. We additionally computed the distance $d$, between the origin of the headpack coordinate system and the fitted line (Table 2 ). Both reflect variability in the perch midpoint's trajectories across all trials per bird, which could be due to the bird fixating on a different point than assumed, eye movements that we are not accounting for, or existing movement of the headpack relative to the head. We used the reference videos of the bird wearing the headpack to inspect whether there could be movement of the headpack with respect to the head, but we found no evidence of this (for further details see Section 11).

To check whether the slope of the fitted line per trial was similar across trials, we computed the orthogonal regression line per trial for one bird (Drogon, Table 3). We computed a deviation metric $\sigma_{\vec{v}_{gaze}}$ for the estimated unit gaze vector per trial, as the square-root of the trace of the covariance matrix:

$$\sigma_{\vec{v}_{gaze}} = \sqrt{\sigma^2_{v_{x,gaze}} + \sigma^2_{v_{y,gaze}} + \sigma^2_{v_{z,gaze}}}, \tag{4}$$

obtaining $\sigma_{\vec{v}_{gaze}} = 0.0463$. The mean gaze vector was $\overline{\vec{v}_{gaze}} = \begin{bmatrix}0.11, & -0.95, & 0.29\end{bmatrix}$, very similar to the gaze direction estimate considering all Drogon trials (Table 2). We also computed the distance from the origin of the headpack coordinate system to the fitted line per trial, which yielded a mean value of $\overline{d} = 65.56$ mm and a standard deviation of $\sigma_d = 22.88$ mm, again showing the variability in the landing perch midpoint trajectories per trial.

**Normal to sagittal plane estimate.** The results of the normal to sagittal plane fit, for all trials per bird, are shown in Table 4. In this case the error in the RMSE metric is the residual of the least-squares problem described in the main text. The samples are the frames in mid-flight phase for all trials per bird. Figure 6 shows the estimated direction $\vec{n}_{sagittal}$ for the Drogon trials.

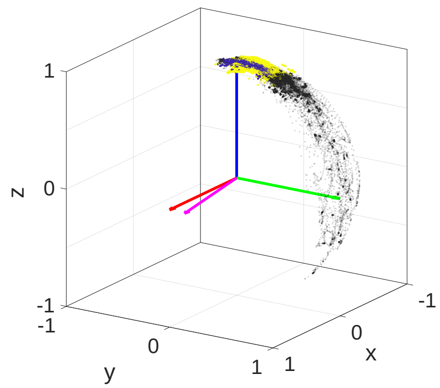


Figure 6: **Estimation of normal to the sagittal plane.** We estimated the normal to the sagittal plane $\vec{n}_{sagittal}$ (magenta), assuming the bird keeps its eyes level during the mid-flight phase. Solid red, green and blue axes correspond to the x, y, z axes of the headpack coordinate system. Dots represent the direction of the z world axis in the headpack coordinate system during the mid-flight phase, for perching (blue) and obstacle trials (yellow). Samples outside of the mid-flight period (black, transparent), show qualitatively larger variation in pitch and roll.

#### 4. Trajectory coordinate system

We define the ***trajectory coordinate system*** as a coordinate system with its y-axis tangent to the forward direction of the trajectory and its x-axis parallel to the horizontal:

$$\vec{y}_T = -\frac{\vec{v}}{||\vec{v}||}, \tag{5}$$

$$\vec{x}_T = \frac{\vec{y}_T \times \vec{z}_{world}}{||\vec{y}_T \times \vec{z}_{world}||}, \tag{6}$$

$$\vec{z}_T = \vec{x}_T \times \vec{y}_T, \tag{7}$$

Table 2: **Results of the gaze direction fits, considering all trials per bird.** The estimated gaze direction $\vec{v}_{gaze}$ and the point in the fitted line $A_{gaze}$ are both expressed in the headpack coordinate system. The distance between the fitted line and the origin of the headpack coordinate system is $d$. RMSE stands for root mean square error, where the error is the distance between the sample points and the fitted line. The number of samples $n$ corresponds to the number of frames in the final approach phase, across all trials per bird.

| Bird | $\vec{v}_{gaze}$ | $A_{gaze}$ (mm) | $d$ (mm) | RMSE (mm) | $n$ |
|---|---|---|---|---|---|
| Drogon | $[0.13, \ -0.94, \ 0.31]$ | $[131.4, \ -969.9, \ 247.0]$ | 68.4 | 65.6 | 1274 |
| Toothless | $[-0.07, \ -0.95, \ 0.31]$ | $[-55.5, \ -969.6, \ 231.1]$ | 82.1 | 46.7 | 847 |
| Charmander | $[0.004, \ -0.97, \ 0.23]$ | $[36.7, \ -1046.4, \ 174.6]$ | 73.9 | 64.7 | 1328 |
| Ruby | $[0.16, \ -0.96, \ 0.22]$ | $[206.5, \ -966.0, \ 158.3]$ | 66.3 | 70.6 | 1353 |

Table 3: **Results of the gaze direction fits per trial (Drogon dataset).** The estimated gaze direction $\vec{v}_{gaze}$ and the point in the fitted line $A_{gaze}$ are both expressed in the headpack coordinate system. The distance between the fitted line and the origin of the headpack coordinate system is $d$. RMSE stands for root mean square error, where the error is the distance between the sample points and the fitted line. The number of samples $n$ corresponds to the number of frames in the final approach phase, for each trial.

| Trial | $\vec{v}_{gaze}$ | $A_{gaze}$ (mm) | $d$ (mm) | RMSE | $n$ |
|---|---|---|---|---|---|
| Drogon perching 1 | $[0.10 \ -0.94 \ 0.33]$ | $[136.50 \ -1108.30 \ 306.00]$ | 84.78 | 21.94 | 122 |
| Drogon perching 2 | $[0.13 \ -0.92 \ 0.37]$ | $[112.38 \ -1086.61 \ 317.02]$ | 113.15 | 48.073 | 123 |
| Drogon perching 3 | $[0.17 \ -0.93 \ 0.33]$ | $[212.42 \ -1094.46 \ 310.33]$ | 74.83 | 37.44 | 122 |
| Drogon perching 4 | $[0.12 \ -0.95 \ 0.28]$ | $[122.18 \ -1138.82 \ 285.44]$ | 46.50 | 44.70 | 117 |
| Drogon perching 5 | $[0.13 \ -0.96 \ 0.25]$ | $[196.22 \ -1115.80 \ 263.51]$ | 50.17 | 29.78 | 117 |
| Drogon perching 6 | $[0.12 \ -0.93 \ 0.36]$ | $[149.82 \ -1095.98 \ 315.56]$ | 99.115 | 54.63 | 124 |
| Drogon perching 7 | $[0.09 \ -0.96 \ 0.27]$ | $[172.20 \ -1126.35 \ 248.55]$ | 91.52 | 43.80 | 112 |
| Drogon obstacles 1 | $[0.12 \ -0.96 \ 0.26]$ | $[141.08 \ -686.26 \ 172.10]$ | 59.95 | 62.09 | 57 |
| Drogon obstacles 2 | $[0.07 \ -0.95 \ 0.29]$ | $[65.88 \ -707.29 \ 154.66]$ | 62.08 | 50.43 | 54 |
| Drogon obstacles 3 | $[0.09 \ -0.96 \ 0.28]$ | $[19.18 \ -711.67 \ 158.51]$ | 65.75 | 48.99 | 56 |
| Drogon obstacles 4 | $[0.15 \ -0.96 \ 0.26]$ | $[117.33 \ -699.39 \ 160.86]$ | 29.74 | 61.56 | 58 |
| Drogon obstacles 5 | $[0.11 \ -0.96 \ 0.27]$ | $[85.59 \ -709.25 \ 155.57]$ | 44.30 | 15.51 | 53 |
| Drogon obstacles 6 | $[0.09 \ -0.96 \ 0.27]$ | $[32.18 \ -710.18 \ 156.70]$ | 57.27 | 24.38 | 55 |
| Drogon obstacles 7 | $[0.12 \ -0.96 \ 0.27]$ | $[109.94 \ -697.48 \ 153.10]$ | 50.87 | 73.59 | 55 |
| Drogon obstacles 8 | $[0.12 \ -0.95 \ 0.28]$ | $[83.00 \ -707.49 \ 154.79]$ | 53.41 | 20.28 | 49 |

where $\vec{x}_T, \vec{y}_T, \vec{z}_T$ represent the versors of the trajectory coordinate system, $\vec{v}$ represents the velocity vector of the head, and $\vec{z}_{world}$ represents the world z-axis. We computed the velocity vector of the head $\vec{v}$ with a central differences scheme on the head interpolated trajectory using the *gradient* function in MATLAB.

#### 5. Distance from the origin of the visual coordinate system to the midpoint between the eyes

The origin of the visual coordinate system is defined as the centroid of the headpack markers projected onto the plane defined by the headpack baseplate. The same origin is used for the headpack and the trajectory coordinate system. We estimated how much this point deviates from the midpoint between the eyes for the dataset used in the main text. We used snapshots of the reference video of the bird wearing the headpack (Figures 7 and 8), recorded before the trials.

Figure 7A shows that the origin of the headpack coordinate system is located between the segment connecting markers 2 and 4. In Figure 7B this segment is highlighted, along with the line connecting both eyes and in Figure 7C the approximate direction of the sagittal plane is shown (magenta). From these we assume that the distance between the origin of the headpack coordinate system and the midpoint between the eyes in the direction perpendicular to the

Table 4: **Results of the normal to the sagittal plane fits, considering all trials per bird.** The estimated normal $\vec{n}_{sagittal}$ is expressed in the headpack coordinate system. RMSE stands for root mean square error, where the error is the residual of the least-squares problem solved. The number of samples $n$ is the number of frames in the mid-flight phase across all trials per bird

| Bird | $\vec{n}_{sagittal}$ | RMSE | $n$ |
|---|---|---|---|
| Drogon | $[0.99, \ 0.14, \ -0.01]$ | 0.08 | 5056 |
| Toothless | $[1.00, \ -0.07, \ 0.02]$ | 0.06 | 3543 |
| Charmander | $[1.00, \ 0.01, \ 0.01]$ | 0.03 | 4641 |
| Ruby | $[0.99, \ 0.16, \ -0.02]$ | 0.03 | 5350 |

sagittal plane is smaller than the maximum distance from the origin to marker 2 or 4 (12 mm).

Figure 8 shows the estimates in the direction perpendicular to the headpack baseplate. We selected three frames in which the bird's sagittal plane was almost parallel to the camera plane. We then estimated on each of them the real length of the red segment, based on the known real length of the yellow segment. We obtained a mean value of 17.3 mm.

We therefore estimate the midpoint between the eyes is within 12 mm from the selected origin in the transverse direction (perpendicular to the sagittal plane), and 17.3 mm in the direction perpendicular to the headpack baseplate.

### 6. Visual field map

To map the visual field of the birds onto the rendered output, we used the data reported for Harris' hawks by Potier *et al.* [4]. The authors measured the visual field experimentally, aligning the bird's sagittal plane with a visual perimeter and using an ophthalmoscopic reflex technique. This way they determined the degree of overlap between the retinal margins of the bird's eyes $\Delta\theta$, at several angles measured from the top of its head $\phi$. We digitized the data from the paper (figures 5A, 5C and 6 in [4]) and fitted a cubic smoothing spline with periodic boundary conditions to the relation $\Delta\theta$ vs $\phi$ (RMSE = 1.6°, see Figure 9). From the retinal overlap $\Delta\theta$ we derived the retinal margins for each eye, $\theta_{left}$ and $\theta_{right}$, assuming symmetry with respect to the sagittal plane. We also assumed our estimated gaze direction corresponds to the $\phi$ angle where the binocular overlap is widest ($\phi = 90°$) (in line with the supplementary figure S1 in [4]).

To overlay the visual field map onto the rendered output, we expressed the retinal margins data ($\theta_{left}$ and $\theta_{right}$) in our latitude and longitude coordinates. We defined longitude as the angle between a vector and the meridian defined by the sagittal plane (i.e., the *yz* plane), and latitude as the angle between a vector and the *xy* plane. We mapped the latitude and longitude coordinates of the retinal margins to the corresponding pixel coordinates to identify binocular and blind areas on the equirectangular rendered output.

We used the OpenEXR bindings for Matlab available at https://github.com/skycaptain/openexr-matlab to read the OpenEXR files in MATLAB, and the *tonemap* function to read the HDR images.

### 7. Model of the environment

We used the Python API in Blender 2.79a to define the geometry of the environment, for the two selected trials for rendering. We defined the geometry and locations of the perches and obstacles based on the position of their corresponding markers (placed at the perches' edges and at the centre of the obstacles' tops).

We reduced the perches to their top rungs and modelled them as cylinders of radius 4 cm (based on estimated measures of the actual perches). From reference images we estimated that a line between the centres of the markers on the perch's edges would be approximately tangent to the top rung cylinder (see Figure 10).

We modelled the obstacles as cylinders of 0.3 m diameter. We estimated the offset between the larger markers' centre and the obstacles' tops by computing the mean deviation from 2 m height for these markers. We took this offset into account (14 mm, including the markers' base) when defining the obstacles' dimensions in Blender.

We defined planes for the walls, floor and ceiling based on the scaffolding points derived from the motion capture cameras' positions (see Section 2). For lighting, we added three lamps of type 'Sun' along the longitudinal axis of the room, 5.5 m apart (the middle one with strength 2, the rest with strength 1; we omitted the middle lamp in the black and white renderings). We defined materials with different colours for the perches, obstacles, walls, floor and ceiling, for clarity, and disabled the casting of shadows.

### 8. Rendering of visual input

Because the vergence movements of the eyes were unknown, we rendered the bird's binocular visual input using

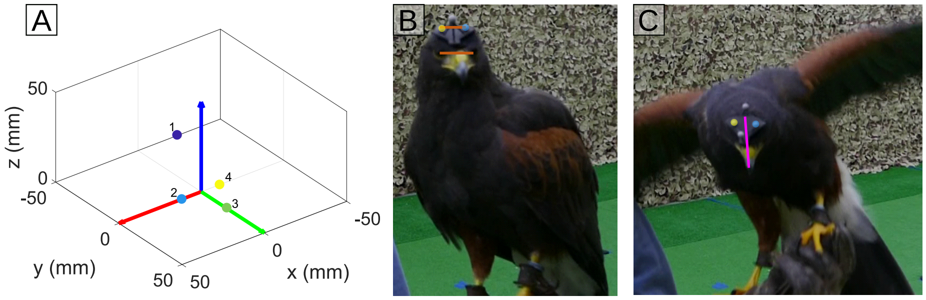


Figure 7: **Distance between selected origin and midpoint between the eyes, in the direction perpendicular to the sagittal plane.** **(A)** Headpack coordinate system and disposition of the markers. The origin (which is the same for the headpack, visual and trajectory coordinate systems) lies approximately between markers 2 (blue) and 4 (yellow). **(B)** The line between markers 2 and 4 (highlighted in blue and yellow respectively) is shown, as well as the approximate line between the eyes. We can see that the midpoint of the line between the eyes falls within the segment connecting markers 2 and 4. We therefore estimate that the distance between the midpoint between the eyes and the origin is at most, the maximum distance between the origin and one of the highlighted markers. **(C)** The magenta line drawn from the bill tip shows the approximate location of the sagittal plane, which contains the midpoint between the eyes.

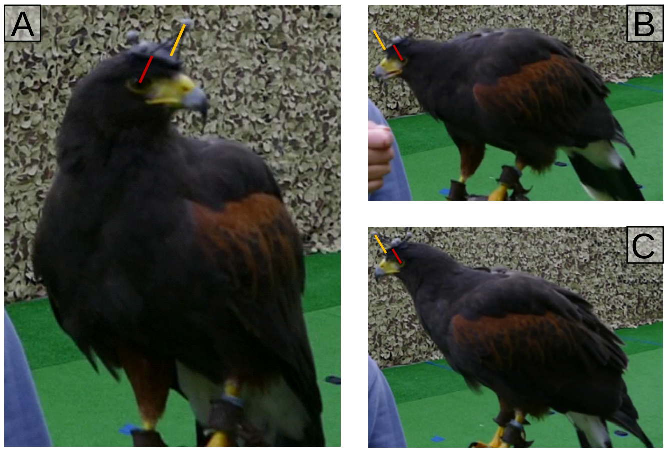


Figure 8: **Distance between selected origin and midpoint between the eyes, in the direction perpendicular to the headpack's plate.** We selected three frames **(A,B,C)** in which the bird's sagittal plane was almost parallel to the camera plane. We then estimated on each of them the real length of the red segment (the approximate distance from the headpack's baseplate to the bird's eye), based on the known length of the yellow segment (the height of marker 1 above the baseplate). We obtained a mean value of 17.3 mm.

a monocular 360° virtual camera with equirectangular projection (latitude ranging from -90° to 90°, and longitude ranging from -180° to 180°). We selected a resolution of 5 pixels per degree latitude and longitude. We animated the camera by setting keyframes for its translation and rotation, which replicated the trajectory of the bird's estimated visual coordinate system, or the trajectory coordinate system. We defined camera keyframes for all frames between the first leg's take-off and the second leg's landing of the trial, and removed any default interpolating animations between keyframes.

We rendered the animations using the Cycles Render engine and GPU compute, in an Dell Inspiron 15-7567 Gaming laptop (Intel Core i7 7700HQ, 16GB RAM, NVIDIA GeForce GTX 1050 Ti 4GB GPU). We output the rendered frames in OpenEXR format, collecting RGB data, depth maps and semantic maps per frame. Every frame took approximately 55 s to render. We rendered two trials of 1661 frames (perching trial) and 1590 frames (obstacles trial).

#### 9. Visual field stabilisation

To investigate the stabilisation of the visual field in flight, we computed heatmaps showing the frequency with which an object appears at each point in the visual field. Figure 11 shows the heatmaps for the second legs of the rendered tri-

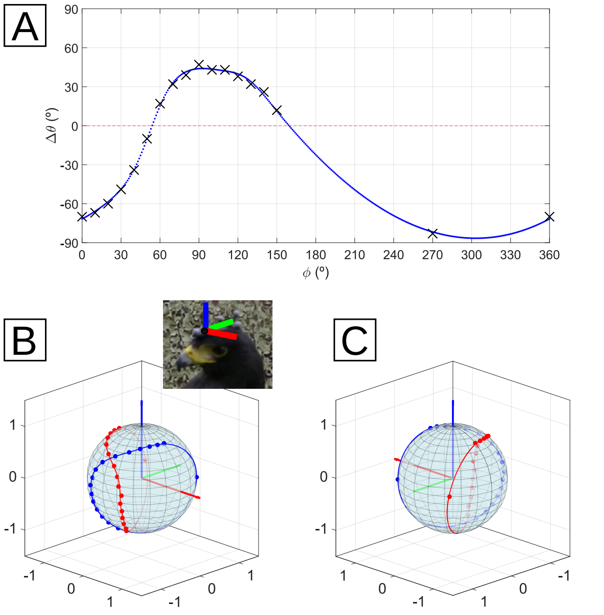


Figure 9: **Visual field of Harris' hawks.** We collected the data describing the visual field of Harris' hawks available in literature [4] and interpolated missing values. **(A)** We digitized the overlap between the retinal margins of the bird's eyes ($\Delta\theta$) at several angles measured from the top of its head ($\phi$). We fitted the data (black crosses) to a cubic smoothing spline (blue dots) of 8 pieces with periodic boundary conditions (RMSE = 1.6°). **(B)** and **(C)** show two views of the retinal margins for the left (blue) and right (red) eye on the unit sphere, in the visual coordinate system. The red, green and blue axes correspond to the x, y, z axes of the visual coordinate system. The inset shows the approximate position of the visual coordinate system relative to the bird's head. The parallels on the sphere are plotted every 9° in latitude and the meridians every 18° in longitude.

als, which were not included in the main text. As in the main text, we excluded from the analysis the 20 frames after each take-off and before each landing, and the frames in which the coordinate system's transform was interpolated (see Section 2). The results are very similar to those for the first legs of the rendered trials given in the main text. The perch is considerably stabilised in the visual coordinate system, compared to the trajectory coordinate system (see Figure 11 A1, A2, B1, B2). The obstacles' edge again seems to be stabilised along the sagittal plane in the visual coordinate system (Figure 11 C1), whereas in the trajectory one they are not within the binocular area (Figure 11 C2). Note that in the second leg of the obstacle avoidance trial the bird takes off from behind the obstacles, so for many frames they appear in the blind area to the back of the bird's head (not visible in the figure).

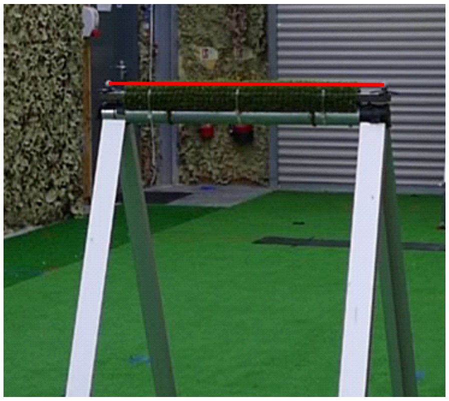

Figure 10: **Position of the markers in the perch.** From reference images like the one shown, we modelled the perch's top rung as a cylinder of radius 4 cm, tangent to the line connecting the markers' centres (red segment).

#### 10. Gaze-shifting behaviour

To investigate the gaze-shifting behaviour, we plotted gaze rays (the gaze vector scaled with the measured depth in that direction) along the head trajectory. We computed the depth in the gaze direction from the rendered depth maps, as the mean depth over the four pixels converging on the gaze direction. Figure 12 shows the gaze rays along the trajectories, for the first and second leg of the two rendered trials (the second leg of the obstacle avoidance trial is the one analysed in the main text, Figure 12 B2). As we did in the main text, we excluded from the analysis the 20 frames after each take-off and before each landing, and the frames in which the coordinate system's transform was interpolated (see Section 2).

The gaze rays for the perching trial (Figure 12 A1 and A2) are shown as side views (i.e., projected on the *yz* plane of the world coordinate system). We did not detect evident gaze shifts from visual inspection in the side view or in the top view. There seems to be a smooth tracking of the landing perch (or nearby points) along the trajectory.

The gaze rays for the obstacle avoidance trial (Figure 12 B1 and B2) are shown as top views (i.e., projected on the *xy* plane of the world coordinate system). We can see some gaze shifts as the bird is making the turn in both cases.

#### 11. Supplementary videos

We include the rendered videos for the two selected trials, for a virtual camera following the visual coordinate system and the trajectory coordinate system. We derived the virtual coordinate system transforms (translation and rotation) from the headpack coordinate system interpolated transforms (see Section 2). We highlighted the frames in which the transform is interpolated with a red contour

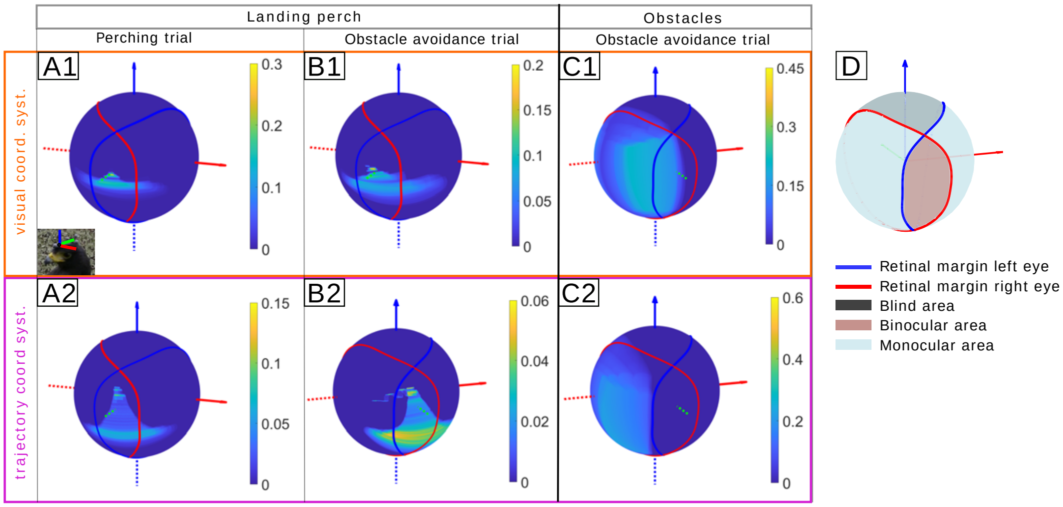


Figure 11: **Visual field stabilisation.** Heatmaps show the frequency with which an object appears at each point in the visual field, throughout a flight. We computed these heatmaps for the second leg of the selected perching and obstacle avoidance trials. Results are shown for a virtual camera following the visual coordinate system **(A1 to C1)**, and the trajectory coordinate system **(A2 to C2)**. As observed in the first leg of these trials, objects are stabilised around the x-axis of the visual coordinate system. The obstacles' right edge is also aligned with the bird's sagittal plane **(C1)**. Inset in A1 shows the approximate orientation of the visual coordinate system relative to the bird's head. Colorbars are scaled based on the rounded-up maximum value observed per heatmap. In C1 and C2 the largest values in the heatmaps occur at the back of the bird's head, in the blind area (not visible in the figure). **(D)** Map of the bird's visual field, derived from Potier *et al.* [4].

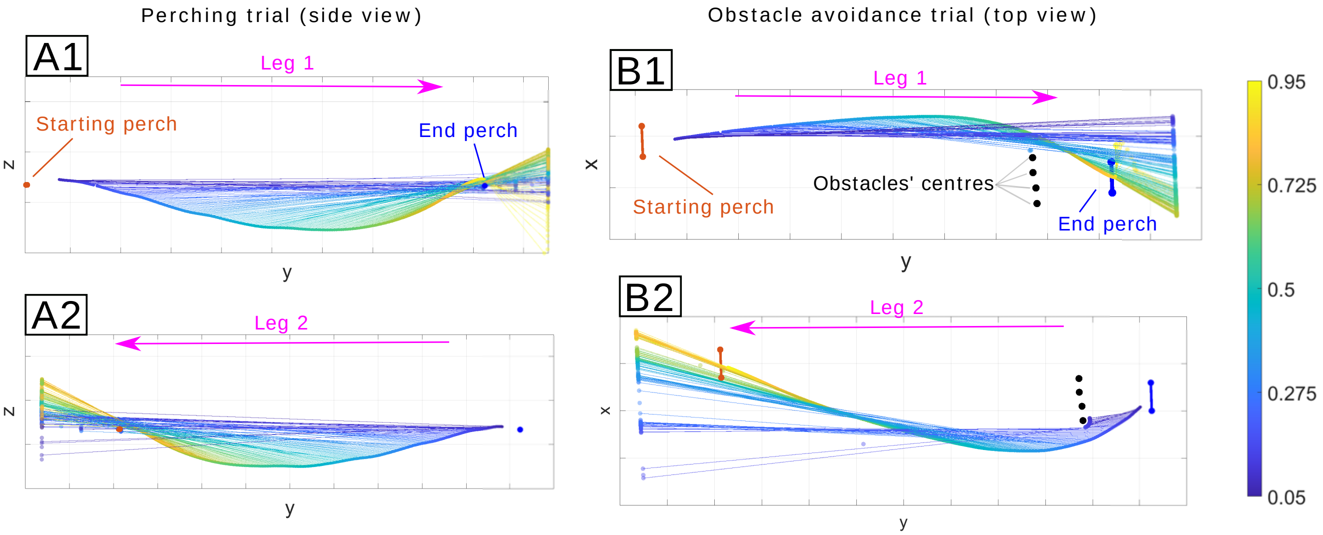


Figure 12: **Gaze shifts in a perching and an obstacle avoidance flight.** Gaze rays are shown every other frame. The colormap indicates time through the trajectory (20 first and last frames excluded). The gaze rays' tips (transparent markers) are plotted for each frame. The magenta arrows at the top of each subplot indicate the direction of flight. For the perching trial, the side view of the bird's trajectory is shown for the first **(A1)** and second **(A2)** legs of the trial. For the obstacle avoidance trial, the top view of the bird's trajectory is shown for the first **(B1)** and second **(B2)** leg trial.

around the rendered image. All rendered videos are played at 20 frames per second (1/10$^{th}$ of the real speed).

Table 5 describes the videos included in the supplementary material. The videos rendered with a virtual camera following the trajectory coordinate system are noisy when the bird is stationary at the perch. This is expected because the tangent of the head trajectory does not vary smoothly when the bird moves its head in a saccadic fashion while

Table 5: **Description of supplementary videos.** All videos are played at 20 frames per second, 1/10 of the real speed. The 'preview' videos were produced with the preview images (jpg format) produced by Blender during the rendering. For the videos with the visual field overlay, we read the high dynamic range images (HDR) from the exr output files and used the *tonemap* function in MATLAB to adjust them for viewing, before overlaying the visual field data. In the videos with the visual field map overlaid, the frames with a red contour correspond to frames in which the position and orientation of the visual coordinate system are interpolated. Note that the videos rendered with a virtual camera following the trajectory coordinate system are noisy when the bird is stationary at the perch. This is expected because the tangent of the head trajectory does not vary smoothly when the bird moves its head in a saccadic fashion while stationary.

| Supplementary videos description |
|---|
| • `Drogon_perching_trial_color_render_preview_20fps`: Color rendering of perching trial, for a virtual camera following the bird's visual coordinate system. |
| • `Drogon_perching_trial_color_render_visual_field_overlay_20fps`: Color rendering of perching trial with visual field map overlaid, for a virtual camera following the bird's visual coordinate system. |
| • `Drogon_obstacle_trial_color_render_preview_20fps`: Color rendering of obstacle avoidance trial, for a virtual camera following the bird's visual coordinate system. |
| • `Drogon_obstacle_trial_color_render_visual_field_overlay_20fps`: Color rendering of obstacle avoidance trial with visual field map overlaid, for a virtual camera following the bird's visual coordinate system. |
| • `Drogon_perching_trial_trajectory_coord_syst_20fps`: Black and white rendering of perching trial, for a virtual camera following the trajectory coordinate system. |
| • `Drogon_obstacle_trial_trajectory_coord_syst_20fps`: Black and white rendering of obstacle avoidance trial, for a virtual camera following the trajectory coordinate system. |

stationary.

In the rendered videos we observed that upon landing, the gaze direction seems to be slightly shifted upwards, off the perch's position. This could indicate how other constraints in the perching manoeuvre (such as avoiding an early stall) become more important than precise vision-based measurements when the landing is imminent. It may also reflect the variability in the gaze direction estimate across different trials. We confirmed that the slope of the gaze direction estimate for each individual trial was very similar to the slope estimated for all trials (see Section 3). However, we found larger variability in the distance to the origin of the visual coordinate system $d$ across trials ($\sigma_d$ = 22.88 mm). This could reflect the bird fixating on slightly different points during approach across trials, or be due to the headpack not being well fixed to the bird's head. To review this, we inspected a reference video of the bird on the falconer's fist, captured before its set of flights. In it, the bird is wearing the headpack and moving its head in a saccadic fashion, and we did not detect any evident movement between headpack and head. The amplitude of the gaze shifts recorded in flight is generally lower than those recorded when the bird is stationary, so we consider the video as a confirmation that there was no significant relative movement between head and headpack in flight.

[3] The Mathworks, Inc., Natick, Massachusetts. *MATLAB version 9.9.0.1538559 (R2020b) Update 3*, 2020. 2
